# A new competitive strategy to unveil the antibiotic-producing Actinobacteria

**DOI:** 10.1101/2023.02.20.529240

**Authors:** Aehtesham Hussain, Umera Patwekar, Anirban Majumder, Aasif Majeed, Yogesh S Shouche

**Affiliations:** NCMR- National Centre for Cell Science (NCCS), Pune- 411007, Maharashtra, India; CSIR- Indian Institute of Integrative Medicine (IIIM), Srinagar, J&K-190005, India

**Keywords:** Microbiome, Actinobacteria, Directed-evolution, NGS, competitive strains, bioactive potential, antimicrobials

## Abstract

The bacterial phylum Actinobacteria encompasses microorganisms with incomparable metabolic versatility and deep resource of medicines. However, the recent decrease in the discovery rate of antibiotics warrants innovative strategies to harness actinobacterial resources for lead discovery. Indeed, microbial culturing efforts measuring the outcomes of specific genera lagged behind the detected microbial potential. Herein, we used a distinct competitive strategy that exploits competitive microbial interactions to accelerate the diversification of strain libraries producing antibiotics. This directed-evolution-based strategy shifted the diversity of Actinobacteria over the experimental time course (0-8 days) and led to the isolation of Actinobacterial strains with distinct antimicrobial spectrum against pathogens. To understand the competitive interactions over experimental time, the metagenomic community sequencing revealed that actinobacterial members from families *Nocardiaceae* and *Cellulomonadaceae* with relatively increased abundances towards end, are thus competitively advantageous. Whilst comparing the Actinobacteria retrieved in the competitive strategy to that of the routinely used isolation method, the Actinobacteria genera identified from competitive communities differed relatively in abundances as well as antimicrobial spectrum compared to actinobacterial strains retrieved in classical method. In sum, we present a strategy that influences microbial interactions to accelerate the likelihood of potential actinobacterial strains with antimicrobial potencies.

## INTRODUCTION

Actinobacteria are one of the diverse groups of microorganisms examined over decades for their biosynthetic abilities. Two-thirds of the clinically successful antibiotics along with a plethora of other drugs and specialized molecules are tracked to actinobacterial origin [1–4]. The recent advent in genome sequencing enabled visualizing these versatile bioactive producers and unveiled diverse biosynthetic gene clusters encoding distinct classes of molecules in their genomes whose chemical compounds have not been identified [5–8], which renders these microorganisms an attractive reservoir to access their potential for new antimicrobials [4].

The increasing antibiotic resistance and challenging treatment options for multidrug-resistant infections [9,10] along with a dramatic decline in the discovery rates of novel classes of antibiotics [11,12,13] create antibiotic crisis. Together, it necessitates the urgency for developing innovative strategies at the early stages of the antibiotic discovery engine. To mine the actinobacterial reserves, designing innovative strategies for isolating potential actinobacterial strains may improve the likelihood of novel antibiotic producers. Nevertheless, early-stage diversification of strain libraries using samples rich in bioactive microbes or explicitly targeting the productive genera may reduce the chances of isolating the common producers.

Though over time, distinct methods have been developed to selectively isolate Actinobacteria that include supplementation of media with antibiotics, pre-treatment of samples, or providing enrichments [14–18]. However, accessing the Actinobacteria resources by generalized approaches, developing new drugs is becoming more difficult [19,20], and using these classical/common approaches, the chance of getting new antibiotic producers in randomly picked Actinobacteria has been estimated very less [21,22]. Nonetheless, utilizing the traditional techniques and the replication (re-isolation of known molecules) remain the major problems in harnessing actinobacterial sources.

Since, the microorganisms through the process of natural selection have adapted to produce defensins or attractants that provide them with an evolutionary advantage over their competitors [23, 24]. This adaptation occurs quickly in rapidly dividing microorganisms under tough selection pressure. As, peptides and small-molecule antibiotics are significant mediators of resource competition and persistence in a competitive environment [25, 26], microbial interactions act as the triggers and cues to activate gene clusters [6, 27–30]. To increase the discovery rates of potential Actinobacteria, we used a novel competition-based strategy to provide a window for the selective establishment of potential antibiotic-producing bacteria, allowing resources to encourage the interference competition among the microbial species for establishing antibiotic-producers, that may be subsequently used to discover the potent strains which may be exploited for revealing the biosynthetic pathways or discovering new bioactive molecules. While competition-based isolation seems appropriate for the establishment of more protective microbiomes [31–33]. However, the production of bioactive molecules is a complex phenomenon, which leads to an array of chemical structures and resistance mechanisms. The correlation between the characteristics of antibiotic production and the resistance phenomenon is expectedly governed by natural selection [34–36]. Our design exploits the natural selection of more competitive strains along with enrichment conditions in liquid suspensions that also influence the retrieval of actinobacterial cultures. Consequently, non-producing bacteria likely have growth rates faster than antibiotic-producing microorganisms and are often more common in the environment than producing microorganisms as these organisms do not have a load of antibiotic production [37–38]. In the competition approach, where conditions of antibiotics as selection pressure may also favour the vertical transmission of primary antibiotic producers among interacting communities that may cause the selective horizontal recruitment of extra antibiotic-producing microorganisms in-vitro.

Here, we used some traits of actinobacterial competition in an up-front strategy to increase the possibility of more competitive actinobacterial strains as a result of interactions among Actinobacteria. We also study the dynamics of the actinomycetes during competition and screened actinobacterial isolates against target pathogens.

## RESULTS

### Strategy development and dynamics of Actinobacteria in the competition

To investigate a new strategy aimed at isolating the antimicrobial-producing Actinobacteria, we used soil samples collected from North-western Himalayas. In addition to isolating monocultures by the commonly used (standard dilution pour plate) method, Actinobacterial isolation was carried out by setting a new competitive procedure using distinct media types (**Figure 1a**). In the competition strategy, we used selective pressure for organisms with better adaptability to reveal competitive actinobacterial producers from naturally enriched mixed microbial populations within the sample. In the competition method, CFUs of the bacteria increased by ~ 3 log folds over 8 days and, we observed dramatic increase in recovery of Actinobacteria from sample suspensions incubated in the 7 different media types. Overall, the bacterial CFUs were found to increase over the eight-day time course. From **Figure 1b,** it is obvious that over 8 days (0-8 days), CFU counts consistently increased in all media types. A greater rate of microbial CFU increase was observed within day2 and day4. However, the majority of Actinobacteria were recovered between the 4th and 8th day of the experimental time course.

**Figure 1.**
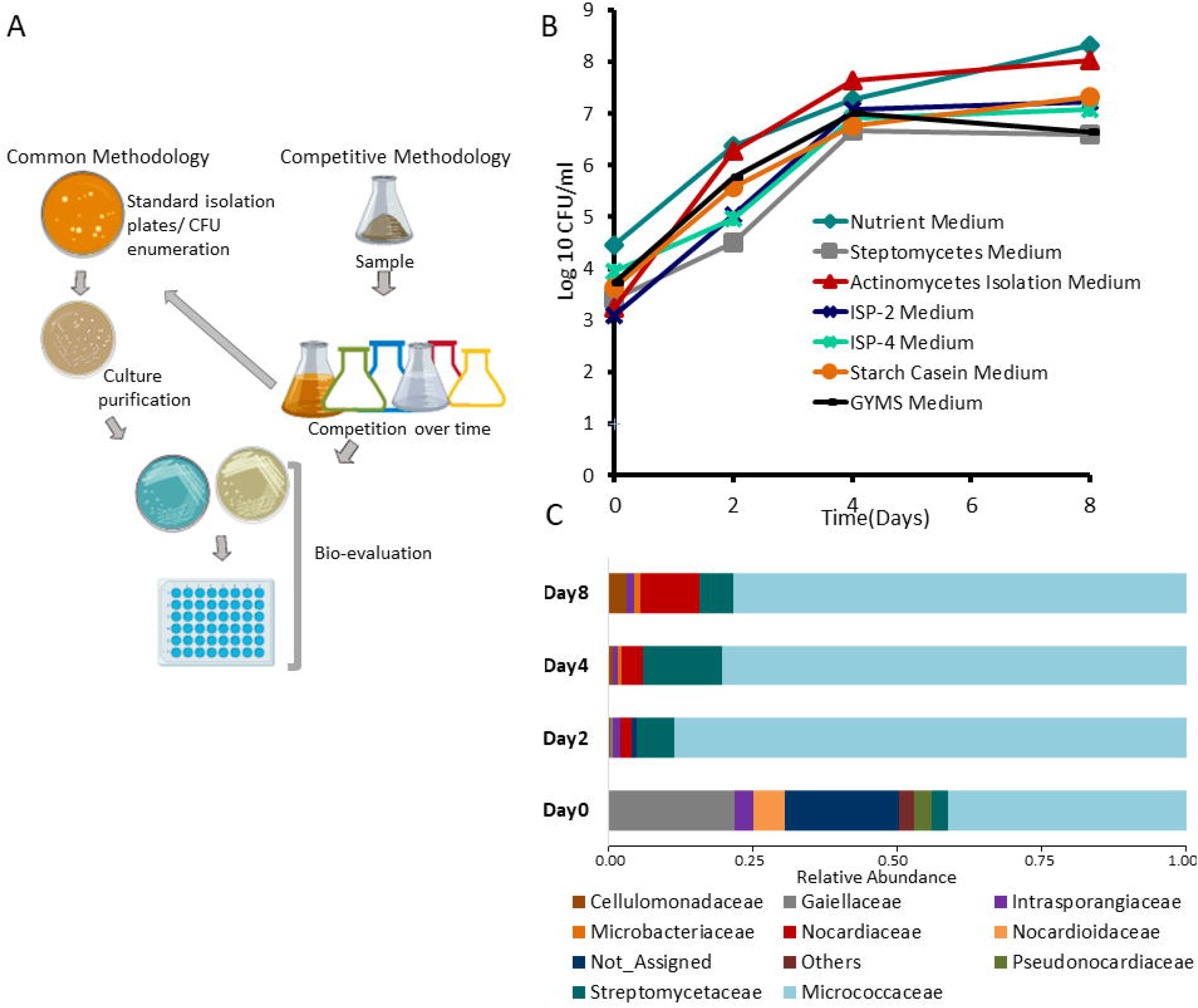
Experimental design and strategy development. **(a)** Experimental design showing an overview of developmental events during the study. **(b)** The fold change of CFUs of bacteria during the study. Each microbial CFUs were recorded as a combination of media and time point. The composite samples were cultured for 8 days for enriching competitive bacteria. At intervals of 0, 2, 4, and 8 days, quantitative CFUs were performed on each media type. **(c)** The relative abundance of actinobacterial families at specific times to the combination of all media types.

We sought to understand the dynamics of the community assembly and recognize the changes that occurred across the eight days of experimental competitive succession using NGS. Using DADA2 1.8 Pipeline, a total of 807126 high-quality (>Q30) reads were recovered after filtering from 2387005 input reads acquired from sequencing. Sequences were assigned to 5610 ASUs.

Assigning the taxonomic affiliations revealed that microbial community changed dramatically over eight days in competitive experiments in comparison to starting day (reflecting original members within the samples). The analyses revealed that members of the phylum Proteobacteria were relatively higher. The microbial members from Proteobacteria, Actinobacteriota, Firmicutes, Acidobacteriota, Verrucomicrobiota, Bacteroidota, Chloroflexi and Methylomirabilota were found relatively abundant in initial experiments. However, there was a significant reduction in relative abundances of phylum Acidobacteriota, Verrucomicrobiota, Chloroflexi and Methylomirabilota over time, but members of phylum Proteobacteria, Actinobacteriota, Firmicutes, Bacteroidota were dominating (**Supplementary Figure S1**). The changes in the community diversity also accompany the significant changes in the family-level composition and actinobacterial increase over time was considerable.

Since the phylum Actinobacteria remain the major producer of bioactive and competitive strains, we extracted 82523 Actinobacteria-affiliated 16S rRNA reads to determine their distribution. A detailed analysis showed that the phylum Actinobacteria was represented mainly by members of orders Micrococcales, Corynebacteriales, Gaiellales, Streptomycetales, Propionibacteriales, Pseudonocardiales and Solirubrobacterales (families: *Micrococcaceae, Gaiellaceae, Nocardioidaceae, Streptomycetaceae and Intrasporangiaceae*). However, Micrococales members were the most abundant class of Actinobacteria over the competitive course. The members of *Micrococcaceae* and Streptomycetaceae increased by Day4 and members of the *Gaiellaceae* reduced significantly (**Figure 1c**) as the least bioactive potential bioactive is observed from members of Gaiellaceae. Towards Day8 of the competition, there was a reduction in *Micrococcaceae*, but a significant shift was noted towards the *Nocardiaceae* and *Cellulomonadaceae* **(Figure 2a**), which is represented by an increase in members of these families and is suggestive that bacteria in *Nocardiaceae* and *Cellulomonadaceae* may be the resource of more competitive bacteria.

**Figure 2.**
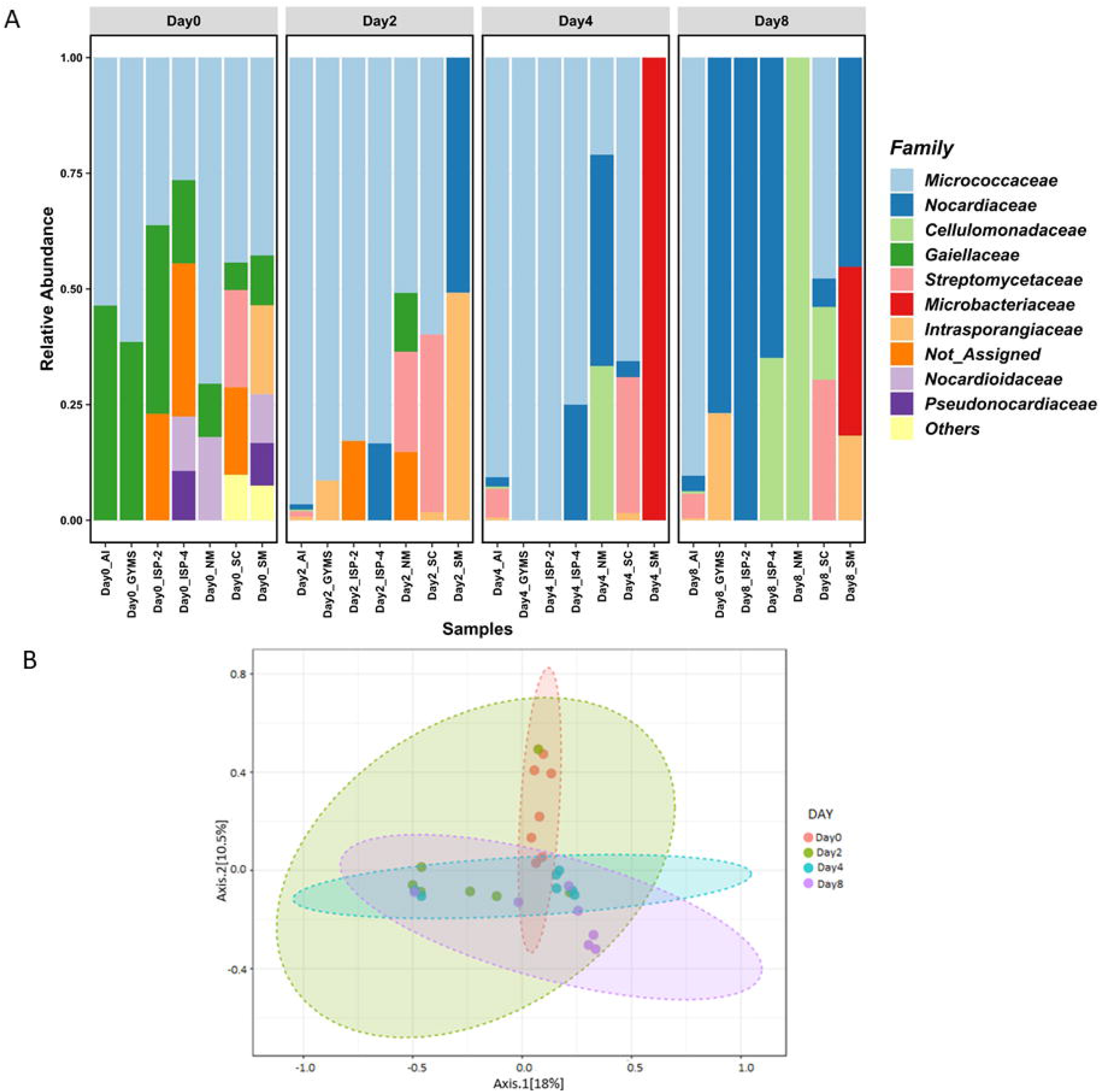
Phylogenetic dissimilarity of microbiota in the experimental period. **(a)** The relative abundance of actinobacterial families for each pairwise combination of time point and media type. The fold enrichment of each of the top nine most abundant actinobacterial families for each time point media type: Nutrient Medium (NM), Streptomyces Medium (SM), Actinomycetes Isolation Medium (AI), Yeast malt media (ISP-2), Inorganic Salt Starch Medium (ISP-4), Starch Casein Medium (SC), and GYMS Medium-agar. **(b)** The principal coordinate plot of all samples is generated using the Bray–Curtis distance; samples are coloured for each combination of sample type and time.

To investigate the changes in relative abundance patterns associated with media types, we analysed the average taxonomic signatures distributed over the 7 different media types, and found the distinct influence of media over actinobacterial distribution (**Supplementary Figure S2**). The Nutrient Medium (NM) substantially favored the growth of Streptomycetaceae which later shifted to *Cellulomonadaceae* on day8. In Actinomycetes Isolation Medium (AI), the relative abundance of the *Micrococcaceae* was higher, but the relative abundance of *Streptomycetaceae* and *Nocardiaceae* significantly increased over time in the competition. In addition to *Micrococcaceae*, Starch Casein Medium (SC) supported the growth of *Streptomycetaceae* in all days in competition. In Streptomyces Medium (SM), the relative abundances of the actinobacterial community shifted towards *Microbacteriaceae*. However, throughout the experimental conditions, we observed the populations of non-assigned/ candidatus taxa decreased overall in all of the media, which is suggestive of designing the special conditions for their enrichments.

The increasing numbers of Proteobacteria may be attributed majorly to genus *Acinetobacter* (**Supplementary Figures S3.a,b**), which has been categorized as highly resistant to present antibiotic drugs. However, over time, we have observed a significant reduction of *Acinetobacter* populations, hinting towards the possibility of enrichment of strains producing new antimicrobials against such resistant microorganisms. Principle Coordinate Analysis (PCoA) of different media-conditioned samples across the time showed a distinctive parting among the bacterial species at different time points (**Fig. 2b**). The directed evolution strategy increased the diversity among the microbial communities across day2 in various media. Comparatively, microbiomes from day4 and day8 samples were varied against day0.

### Cultivation and identification of Actinobacteria

Next, we set out to identify the axenic actinobacterial cultures isolated over key time points under standard laboratory-rearing conditions. A total of 443 presumptive Actinobacteria were isolated in this study using both classical procedure and competition-based strategy with the media and procedures. After dereplication by morphological characteristics, 120 morphologically distinct actinobacterial strains were retained further for study. The closest type strains are shown in **Supplemental Table S2.** The actinobacterial monocultures were identified using MALDI-TOF/MS-based identification (**Supplementary Table S3**) and 16S rRNA gene sequencing-based identification (**Supplementary Table S4).** However, using MALDI-TOF/MS only 10% of the isolates showed reliable identification owing to the limited library of spectra, and thus were identified by 16S rRNA gene sequencing. Together, MALDI-TOF/MS and 16S rRNA gene sequencing-based identification showed that actinobacterial morphotypes represent **68 different species** belonging to **17 different actinobacterial genera** viz. *Streptomyces, Amycolatopsis, Paenarthrobacter, Arthrobacter, Rhodococcus, Nocardia, Brachybacterium Kocuria, Microbacterium, Brevibacterium, Corynebacterium, Agromyces, Micromonospora, Gordonia, Nocardiopsis, Cellulosimicrobium*, and *Micrococcus*. Using the common methodology, the actinobacterial members from genera: *Streptomyces, Amycolatopsis, Paenarthrobacter, Arthrobacter, Rhodococcus, Nocardia, Brachybacterium, Kocuria, Microbacterium, Brevibacterium*, and *Corynebacterium* were isolated. The genus-wise distribution of isolated strains revealed that competition strategy could cover 16 (94.1%) genera of 17 different actinobacterial genera isolated in the study. However, using the classical methodology, only 11 (64.7%) of the genera were represented (**Fig. 3a**). The additional actinobacterial members from *Gordonia, Nocardiopsis, Agromyces, Micrococcus, Micromonospora, Cellulosimicrobium* were retrieved from competition strategy. However, *Brachybacterium* was represented in classical method only (**Figure 3b, supplementary Table S5).** Comparatively, the competition strategy also lead to increased diversity of actinobacterial genera (**Figure 3b, c**). Based on the closest hit, species-wise, we found 38 (55.9%) different species of the total species in the commonly used approach while 41 (60%) species of a total of 68 different species were represented in the competition method (**Supplementary figure 4**). Besides isolating diverse actinobacterial isolates, many putative novel species with 16S rRNA gene sequence similarity </= 98.7% were retrieved which warrants detailed characterization for species delineation. The competition method resulted in cultivation of additional actinobacterial genera and even yielded putative novel *Nocardia* species.

**Figure 3.**
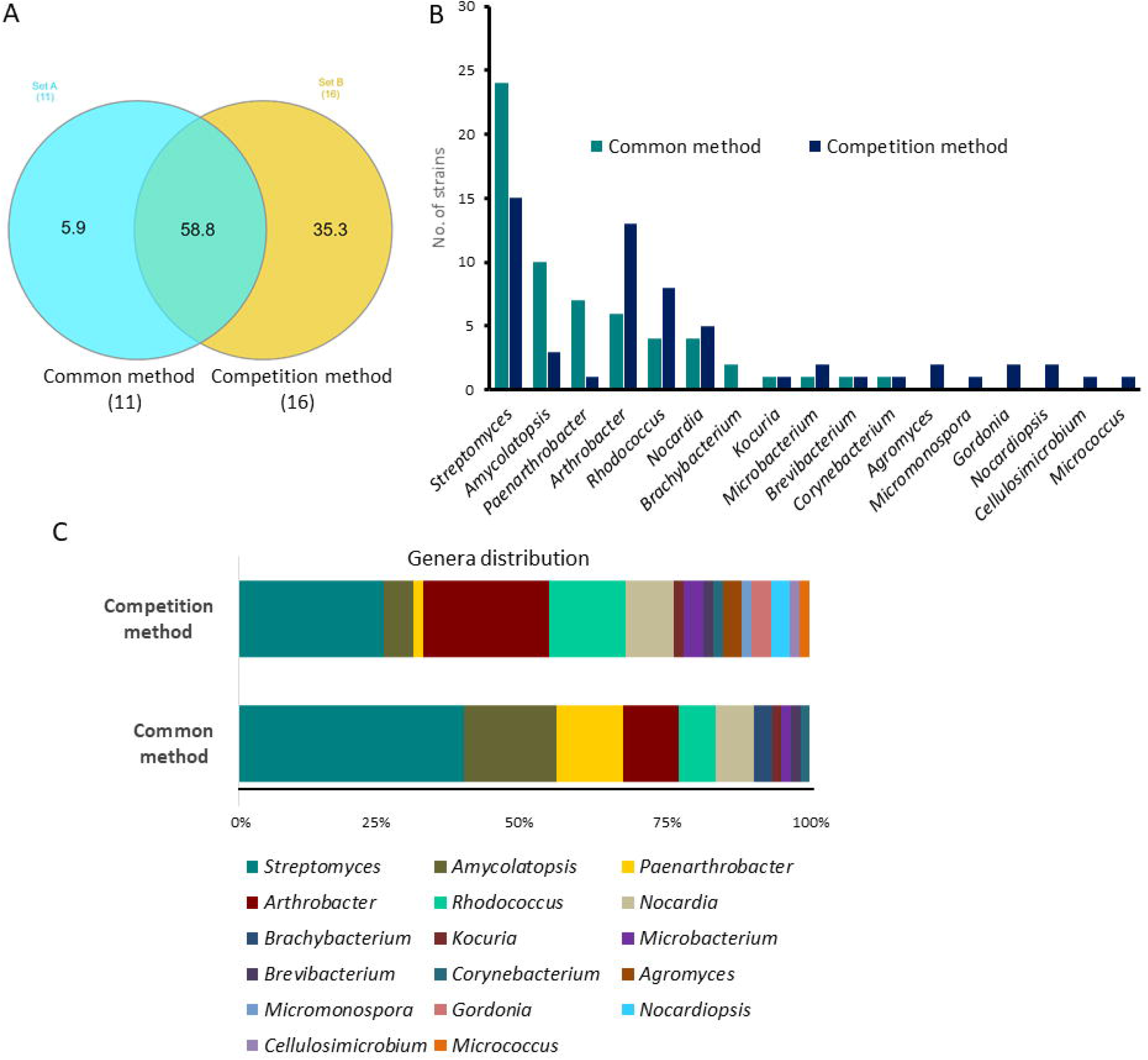
Actinobacterial distribution in normal and competitive procedures. **(a)** The figure (Venn diagram) shows unique and shared genera between the methodologies. Set A represents total genera in the normal method (n=11); set B represents genera in the competitive method (n=16). The genera unique in the normal method (n=1); the genera unique in the competitive method (n=6); genera common in both methods n=10. **(b)** Bar representation showing the number of actinobacterial genera retrieved by common and normal and competition procedures. **(c)** The comparative actinobacterial genera level distribution in normal and competitive procedures.

### Antagonism and bioactive potential

Also, one of our goals was to move beyond the descriptive taxonomy measures to understand the potencies of the microbial populations subsequently retrieved over time in a competitive course and compare their antimicrobial potencies.

All the strains were evaluated against a pathogenic panel including standard laboratory strains including few microorganisms from ESKAPE list of microorganisms viz Gram Positive (*Staphylococcus aureus, Micrococcus luteus, Enterococcus faecalis*) and Gram-negative (*Escherichia coli, Pseudomonas aeruginosa, Klebsiella pneumoniae, Acinetobacter baumannii*). In 50 potential inhibitory actinobacterial strains, 39% (24 strains) of the Actinobacteria retrieved in the conventional method and 44% (26 strains) amongst actinobacterial strains isolated in the competition method inhibited the growth of pathogenic microorganisms, as determined by the antagonism test. Based on the closest hit, the bioactive actinobacterial strains were affiliated with the genera *Streptomyces* (54%), *Amycolatopsis* (16%), *Rhodococcus* (8%) and, 4% each from *Arthrobacter, Brachybacterium, Brevibacterium, Nocardia, Paenarthrobacter* in classical methodology. However, in competition strategy, the potential inhibitory genera were found as *Streptomyces* (26%), *Arthrobacter* (15%), *Rhodococcus* (15%), *Nocardia* (11%), *Nocardiopsis* (7%) and 3.8% from each *Agromyces, Amycolatopsis, Cellulosimicrobium, Gordonia Kocuria, and Paenarthrobacter*. Though, the bacterial inhibitions revealed members from the actinobacterial genus *Streptomyces* were dominant in both methods and accounted for 54% and 26% of active strains in normal or competitive methods respectively **(Figure 4a)**. However, in the competitive method, Actinobacteria from genera *Arthrobacter, Rhodococcus, Nocardia, and Nocardiopsis* displayed higher inhibitory potential. Comparatively, the strains isolated from the competitive method showed high representations in the inhibition assay (**Figure 4b)**. The strains isolated towards the end of competitive experiments were following the NGS amplicon sequencing data. The antimicrobial bioactive actinobacteria from genus *Nocardia* and *Amcolamycotais* isolated on day8 of the competition genus were antagonistic against the Acinetobacter strains. We isolated potential strains from genus *Nocradia* and *Nocraiopsis* towards day8 of competitive experiments that were in accordance with NGS amplicon sequencing data revealing that families *Nocardiaceae* and *Cellumonadie* are competitively dominating at day8. Nevertheless, it was observed that populations of the microbes towards the end of experiments were antagonistic against the *Acinetobacter baumannii* strains showing these producers may have chemical scaffolds to target such resistant bacteria. A closer assessment of inhibitory strains revealed that actinobacteria isolated from the common method were least active against *Acinetobacter Baumannii* in comparison to bacteria isolated in competition experiments. In addition, in the course of the competition experiments, we investigated changes in relative abundance patterns of all bacteria and the striking change observed was that resistant non-actinobacterial members *Acinetobacter spp*. and *Stenotrophomonas spp*. were increasing among non-actinobacteria (**Supplementary Fig. 3b**) that were also identified during cultivation of microorganisms showing bioactivity against *A. baumannii*.

**Figure 4.**
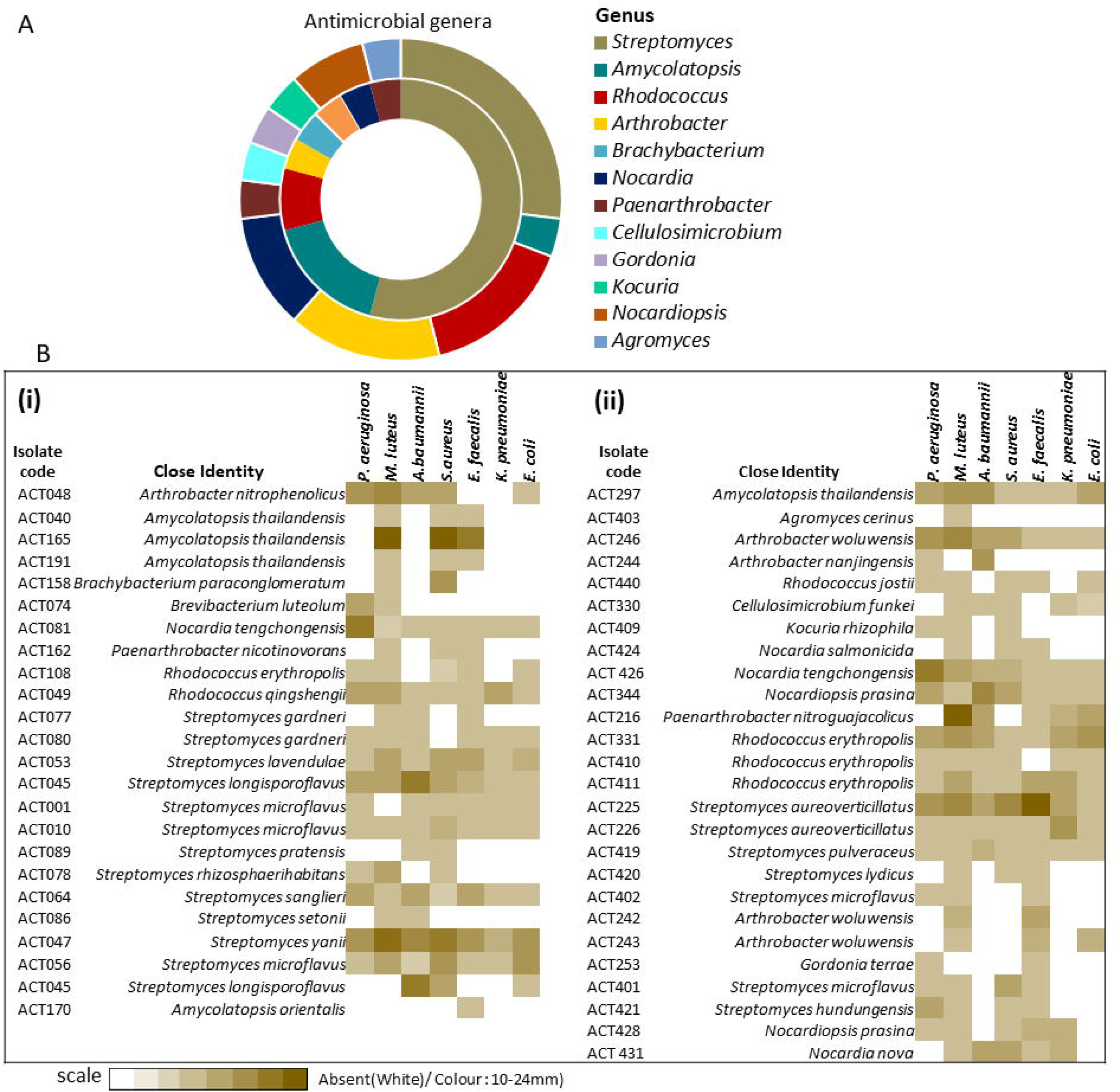
Antimicrobial spectrum of Actinobacteria in two strategies. **(a)** The comparative antagonistic bioactive actinobacterial genera in classical (common) and competition strategies. A figure shows the comparative genus-level distribution of antimicrobial actinobacteria retrieved by both methodologies. The inner ring represents antimicrobial actinobacterial genera retrieved by common method and the outer ring represents antimicrobial actinobacteria retrieved by competition procedures. **(b)** Heat map representation of the antimicrobial spectrum of potential actinobacteria isolated with classical (i) and competition (ii) methodologies. The pathogenic panel includes the pathogenic microorganisms *Staphylococcus aureus, Micrococcus luteus, Enterococcus faecalis, Escherichia coli, Pseudomonas aeruginosa, Klebsiella pneumoniae*, and *Acinetobacter baumannii*.

Thus we identified an increased activity spectrum against test pathogens and found dramatic changes in bacteria associated with antibacterial activities. The study also demonstrated that competition-based directed evolution can be used in detecting the potential strains with enhanced potency.

## DISCUSSION

Actinobacteria have enormous natural product variability between strains [1], and only the genus *Streptomyces* is expected to produce more than 100,000 antimicrobials [5, 39]. Owing to such biosynthetic novelty, innovative strategies are important to surface the potential Actinobacterial strains.

The classical bacterial taxonomy-guided discovery workflow will continue to access microorganisms, however, mining the actinobacterial pools requires innovative strategies to surface strains with better efficacy against the plethora of drug-resistant microorganisms. For taping biosynthetic potential of microbial resources, multiple strategies including genome mining approaches [40,41] metagenome-sequencing-based approaches [42], chemical profiling-based approaches [43,44], enrichment based on intrinsic antimicrobial resistance for likely producers [45], or elicitation to activate silent gene clusters [46] have been developed. Each of these strategies has its distinctive benefits, however, most of these strategies involve exhaustive genetic manipulation, chemical analysis, and high-throughput culturing for discovering biosynthetic potential. Despite the advances in sequencing approaches, the screening procedures are largely allied to microorganism cultures [47]. Nevertheless, various studies reported many discrepancies in chemical features in microorganisms despite belonging to the same taxonomy based on 16S rRNA gene sequences in bacteria from distant origins [48–50]. Thus, bioprospecting microorganisms just by identifications may not be connected correctly to biosynthetic potential.

Since microorganisms have adapted through the process of natural selection to synthesize molecules that afford evolutionary advantage [23, 24]. Evolution in nature relies on the application of selection pressure across a population and thus may select for distinct structural properties or molecular targets or natural products to mediate interaction with other organisms, thus drives the development of bioactive molecules [51]. Ecological diversification seems a rigorous rationale to detect new biosynthetic gene clusters from environmental microbes which can maximize the chemical novelty. We attempted to mimic the directed-evolutionary processes that allows to introduce variability across the microbial population. Under selection pressure, more successful microbes have a chance to survive and propagate for the next round of selection. Here we, harness the speedy directed evolutionary processes to surface the strains with better bioactivity owing to their natural product biosynthesis. So, rather than a taxonomy-guided approach, this approach relies more on competitive interactions, providing a new opportunity to directly harness the microbes having desired bioactivities.

The interspecies interactions that influence microbes or their secreted chemical cues can be employed as inspiration to harness the microbial strains producing new molecules [52,53]. These interactions are mediated by competition for resources, exchange of chemical cues, or other mechanisms. Antibiotic production is more likely induced by cues from closely related strains which share secondary metabolite biosynthetic genes [54]. We followed a completion-based strategy that relies on specific microbial interactions involving competition and adaptation to maximize the possibility of likely antibiotic producers. This strategy dramatically shifted the diversity of producing actinobacteria with different genera compared to the original communities. The changes in the actinobacterial populations may be attributes due to microbes with greater survival and adaptability of likely producers. The Actinobacteria retrieved during the competitive strategy inhibit microbial growth of microorganisms suggesting that likely antibiotic producers may yield molecules with a different mechanism of action. On the applied front, one of the attributes of such competition strategy may be also viewed as analogous to the community genetic information hypothesis to design medicinal chemistry.

Since evolutionary processes are mediated through traits encoded by genomes of organisms and antibiotic production has been found strain-specific trait [55]. Using the competition method, we retrieved actinobacterial strains from genera *Streptomyces, Arthrobacter, Rhodococcus, Nocardia, Nocardiopsis* and *Agromyces, Amycolatopsis, Cellulosimicrobium, Gordonia, Kocuria, and Paenarthrobacter*. Also, beyond the descriptive taxonomy measures we subsequently attempted to understand the potencies of isolated microbes during the competition course. An increased activity spectrum of strains was observed during the experimental time course. Knowledge acquired from such experimental shaping of bacterial competition will contribute to an increased understanding of processes important in competition and also to prioritize microbes for discovering next-generation bioactive molecules. The wider antimicrobial spectrum in comparison to strains isolated from normal procedures, however, can also be attributed to the activation of silent biosynthetic gene clusters in competition. Moreover, this enriched competitive approach offers its simplicity, i.e., without using the more labor-intensive methods for retrieving and diversifying the microbe libraries of potential Actinobacteria. Besides, this strategy can complement existing methods designed for isolating the potent Actinobacteria. However, using different liquid growing conditions also influences the retrieval of Actinobacteria members. Thus, strategy highlights the significance of integrating competitive ecological factors into microbial discovery efforts i.e., diversifying the microbes to discover potent antimicrobial strains for augmenting lead discovery.

## MATERIALS AND METHODS

### Strategy development: Actinobacteria Isolation

Multiple soil samples from the North-western Himalayas were taken in sterile sampling bags, transported to the laboratory, and kept at 4°C. To capture as much actinobacterial diversity, we used a range of rich and selective culture media. To isolate Actinobacteria we used both the standard spread plate method (the common method) and a new competitive strategy.

In the standard spread plate isolation method, the samples were serially diluted 10 folds in PBS to 10^-1^ to 10^-7^. Appropriate aliquots from each dilution were spread onto the surface of isolation plates of the different media types: Nutrient Medium (NM)-agar, Streptomyces Medium (SM)–agar, Actinomycetes Isolation Medium (AI)-agar, Yeast malt media (ISP-2)-agar, Inorganic Salt Starch Medium (ISP-4)-agar, Starch Casein Medium (SC)-agar, GYMS Medium-agar (composition: **Supplementary Table S1**) supplemented with Polymyxin B sulphate, Neomycin B 50 μgmL^-1^, Amphotericin B 2.5μg mL^-1^, Cycloheximide 10 μgmL^-1^ to eliminate gram-negative bacteria and fungal growth. Isolation plates were incubated for 14 days at 28°C, and colonies were observed periodically. The majority of actinobacterial isolates were verified by phenotypic examination. The resulting colonies were purified and maintained SC-agar media. The selected strains were stored in 20% glycerol at −20°C.

In a directed-evolution-based competition strategy, we used an enrichment procedure involving seven different media in broth. Briefly, the composite sample was formed and portions of the sample (2gm) were suspended in culture flasks containing seven media types (50 ml broth, **Supplementary Table S1**) that were supplemented with antifungals (cycloheximide 10 μgmL^-1^ and amphotericin 2.5μgmL^-1^) and antibacterials (Polymyxin B 10 μgmL^-1^, Neomycin B 50 μgmL^-1^) to prevent fungal and bacterial growth. To achieve bacterial competition, all flasks containing soil samples were incubated at 28 °C under shaking (100 rpm) for 8 days. 1 mL of aliquots of mixed communities from each of the suspension culture flasks were taken at time points (0, 2, 4, 8 days) for metagenomics community sequencing. Also, 1 ml aliquots from each of suspension flask (7 types) at specified time intervals (0, 2, 4, 8 days) were serially diluted 10 folds in PBS to 10^-1^ to 10^-7^. The dilutions from enriched suspensions were spread onto the respective media agar plates (**Supplementary Table S1**). All the plates were incubated at 28°C for 14 days and looked for different bacterial colonies at regular intervals. The CFU counting was carried out at different intervals for 15 days (day2, day7, day15). New CFUs observed on day15 were added to those already scored earlier. The resulting colonies were purified and maintained respective agar media. The selected strains were stored in 20% glycerol at −80 °C.

### DNA extraction, Illumina library preparation, and 16S rRNA gene amplicon sequencing (NGS)

Total DNA from each time point of experimental microbial communities was extracted from samples using QIAamp DNA mini kit (Qiagen, Germany) following the manufacturer’s instructions, DNA was quantified at 260 nm using NanoDrop One (Thermo Fisher Scientific, Madison, USA) and was quality checked. 16S rRNA gene amplification was performed by targeting V4 hypervariable region using the primer pair (V4 Forward 515F ([5’-GTGCCAGCMGCCGCGGTAA3’] and V4 Reverse 806R [5’GGACTACHVGGGTWTCTAAT3’]) with PCR conditions (at 95°C for 3 min, followed by 25 cycles of 95°C for 30 □s, 55°C for 30 □s, and 72°C for 30 □s; and finally at 72°C for 7 min [56]. The amplified PCR products were purified using AMPure XP beads (Beckman Coulter, Inc., USA). The amplified products were attached with Illumina adaptor sequences and dual indices using the NextraXT library preparation kit (Illumina, USA) by Index PCR. Further, Index PCR products were purified using AMPure XP beads, and library fragment size distribution was checked using TapeStation (Agilent Technologies, USA) and quantified using Qubit 4.0 (Thermo Fisher Scientific, USA). The quantified libraries were diluted to equimolar concentration (4nM), denatured, and finally sequenced with phiX as control using paired-end 2×250-bp chemistry on the Illumina MiSeq 2000 platform (Illumina Inc., USA)

### Microbiome diversity measures and statistical tests

The quality of raw amplicon sequencing data was checked using the FastQC tool [57] and further taxonomy was assigned using DADA2 (1.8) pipeline [58] in R Software 4.1.0. Based on sequence quality plots, the forward and reverse sequences were trimmed to 240bp and 160 bp respectively. The erroneous reads were removed and the forward and reverse reads were merged following the removal of chimeric sequences. After the filtration step, 807126 reads were finally recovered and used for downstream analysis to generate ASVs (amplicon sequence variants) using the sample inference algorithm of the DADA 2 pipeline. The taxonomic assignment was done using the SILVA version 132 database (https://www.arb-silva.de/documentation/release-132/). The final outputs of the analyses were the ASVs count table and TAXA table which were used for further bioinformatics analysis using MicrobiomeAnalyst 2.0 [59]. The output data from the MicrobiomeAnalyst was used for the generation of various graphical plots.

### 16S rRNA gene sequence-based identification of Actinobacteria

Genomic DNA from well-isolated and purified colonies was extracted using cetyltrimethylammonium bromide (CTAB) and phenol-chloroform extraction method as described by Magarvey *et. al*, [60]. DNA was quantified using NanoDrop One (Thermo Fisher Scientific, Madison, USA). The selective amplification of 16S rDNA gene was performed using bacterial universal primers: 27F (5’-AGAGTTTGATCCTGGCTCAG-3’) and 1492R (5’-GGTTACCTTGTTACGACTT-3’), Taq DNA polymerase (Qiagen, Hilden, Germany) in Mastercycler pro S thermocycler (Eppendorf, Germany), with PCR program: Initialization: 95°C, 1 min., following 30 cycles of denaturation: 95°C, 15s; annealing: 55°C, 15s; elongation 72°C, 90s, and final elongation: 72°C, 7 min. The amplification products were purified using the Polyethylene glycol (PEG)□NaCl precipitation procedure [61]. The identity of the Actinobacteria was confirmed by Sanger DNA sequencing with 3730xl Genetic Analyzer (Applied Biosystems) using conserved primers. The generated consensus sequences of the 16S rRNA gene were analysed in the database of validly published bacteria, EzBioCloud [62], and the threshold of 98.7% 16S rRNA gene sequence similarity was first considered as an indication for prospective novel taxa.

### MALDI- TOF/MS-based Identification of Actinobacteria

MALDI-TOF/MS-based identification of bacteria was performed using alpha-cyano-4-hydroxycinnamic acid (HCCA) matrix. The sample preparation and protein extraction was performed as described by Kurli *et. al*, [63]. Samples were analyzed with Autoflex speed MALDI–TOF/TOF mass spectrometer (Bruker Daltonik GmbH, Germany) in linear positive ion extraction mode with mass cut off m/z range: 2000–20,000Da. Mass spectra visualization, spectra-based identification of microorganisms, and dendrogram construction were achieved using MALDI Biotyper software 3.0 (Bruker Daltonik, GmbH Germany). The biotyper score values greater than 2.0 were taken as species-level identity.

### Bioactivity: Antagonism Evaluation

The bioactivity of actinobacterial strains was tested against the pathogens as described by Hussain *et. al*. [64]. The test bacteria were screened in duplicate for the antibiosis potential against pathogenic bacteria (inoculated at density 1X10^5^ CFU) on MHA plates and plates were incubated at 37 °C for 24–48 h. The pathogenic bacteria include lab standard strains: *Micrococcus luteus* ATCC 10240, *Staphylococcus* aureus ATCC 25923, *Escherichia coli* ATCC 25922, *Pseudomonas aeruginosa* ATCC 27853, *Klebsiella pneumonia* ATCC 700603, *Enterococcus faecalis* ATCC 29212 and *Acinetobacter baumannii* ATCC 19606. The inhibition zones were measured in millimeters (mm).

## ACKNOWLEDGMENTS

We gratefully acknowledge the Director NCCS and the Department of Biotechnology (DBT), Govt. of India for support (Grant No. BT/Coord.II/01/03/2016) at NCMR- National Centre for Cell Science (NCCS), India.

## AUTHOR CONTRIBUTIONS

AH designed the study. AH, UP, AM performed experiments. AH, UP, and ANM analysed data. AH wrote the main manuscript. AH and ANM prepared figures. All authors revised the manuscript and approved the final manuscript.

## LEGENDS TO SUPPLEMENTARY FIGURES

**Supplementary Fig S1.** Relative abundances of bacteria (phylum level) in different time points over the study.

**Supplementary Figure S2.** Media influence in competition.

Relative abundance patterns of Actinobacteria associated with different media types. Enrichment is shown independently for each of media type. The soil samples were cultured in competition in enriching media for 8 days. At intervals of 0, 2, 4, and 8 days, quantitative CFU was performed on each media type.

**Supplementary Figure S3.** Microbiota shift (genus level) in the experimental time.

**a)** The relative abundance (genus level) at different time points within competition.

**b)** Relative abundance (genus level) of the top most abundant bacterial genera for each media type and time point.

**Supplementary Figure S4.** Unique and shared species in either methods.

**(a)** Venn diagram shows unique and shared species retrieved by both methodologies. Set

A (left) represents different species retrieved in the common method and set B (right) represents species in the competition method.

**(b)** Actinobacterial species: unique and common retrieved in competition and classical methods. The species mentioned here is based on validly published type strains searched by EzBioCloud or the closely related type strains determined in MALDI Biotyper software 3.0.

