## Supplementary figures and images for "A new competitive strategy to unveil the antibiotic-producing Actinobacteria"

### Supplemental Figure S4

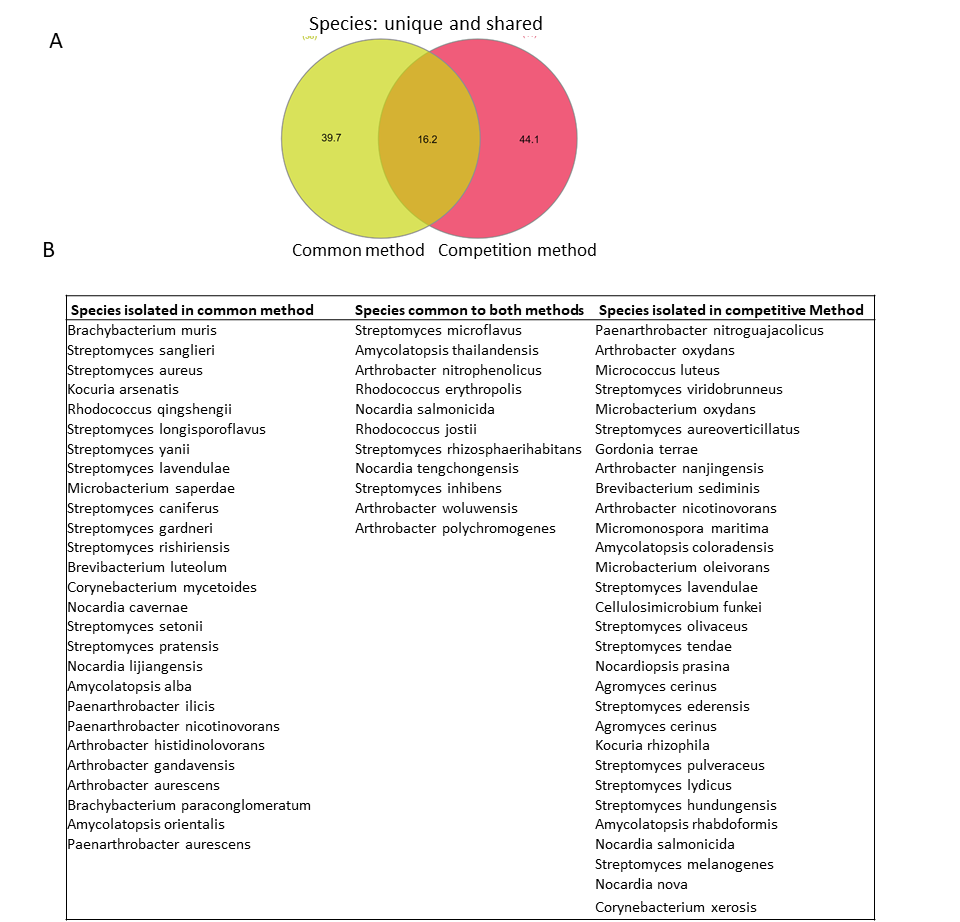

### Supplementary Figure S1

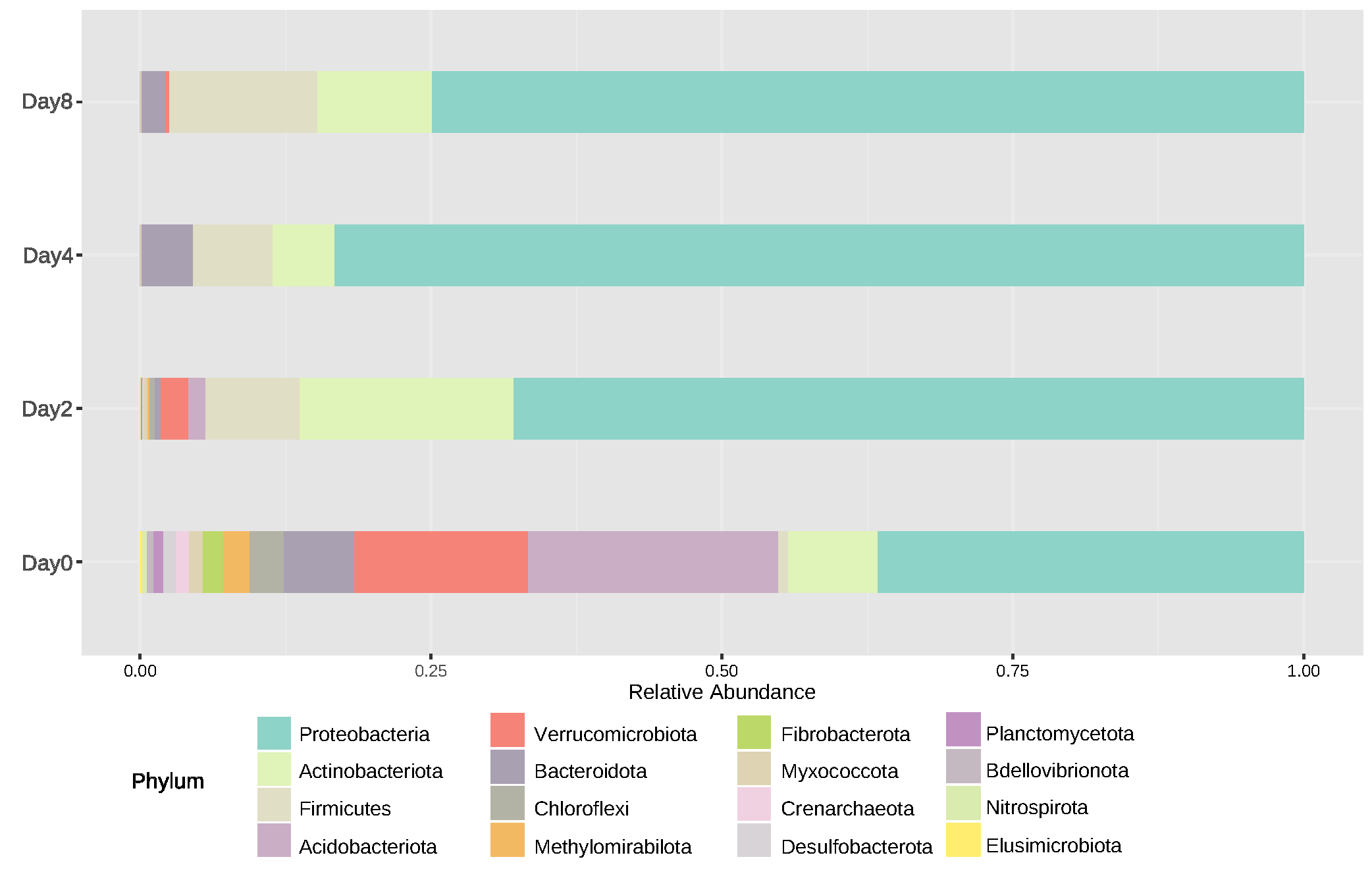

### Supplementary Figure S2

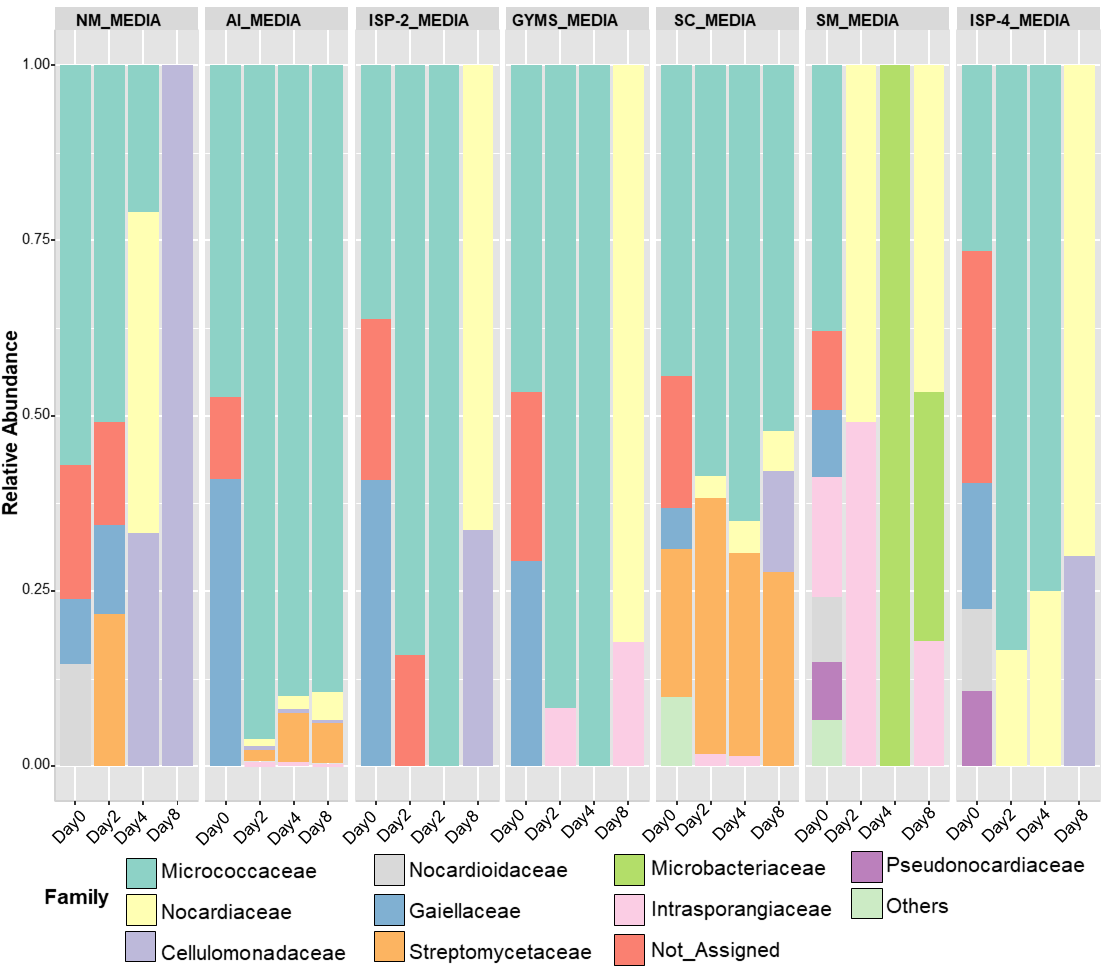

### Supplementary Figure S3

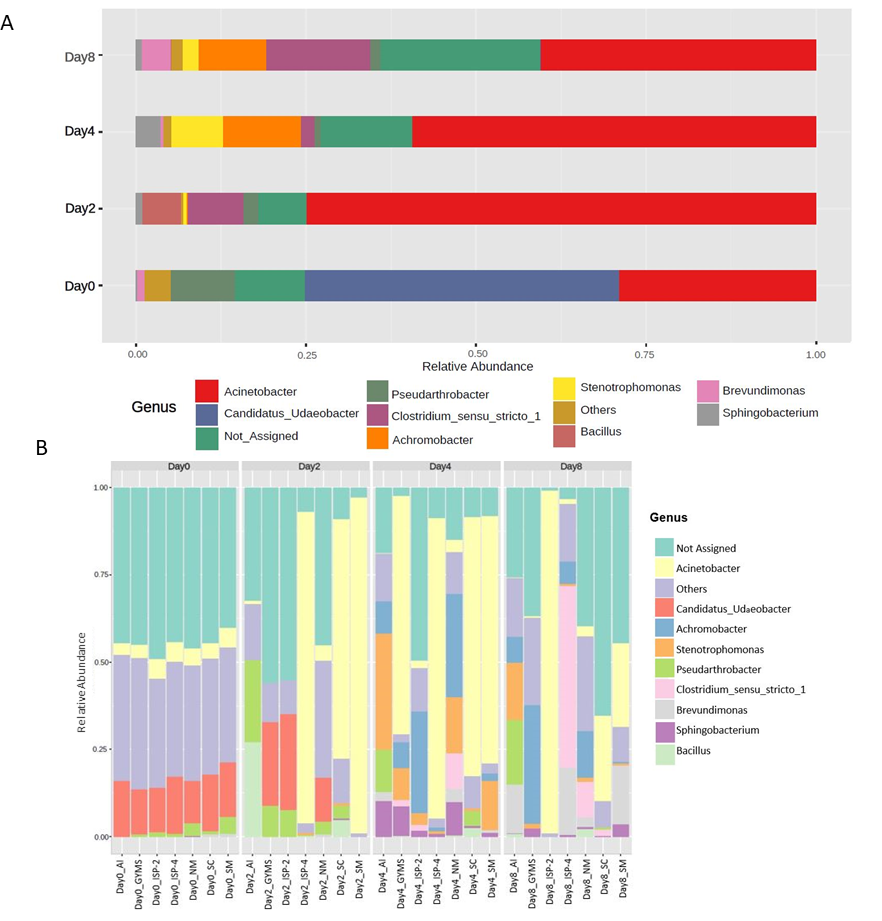
